# Although host-related factors are important for the formation of gut microbiota, environmental factors cannot be ignored

**DOI:** 10.1101/2020.02.03.933317

**Authors:** YeonGyun Jung, Dorsaf Kerfahi, Huy Quang Pham, HyunWoo Son, Jerald Conrad Ibal, Min-Kyu Park, Yeong-Jun Park, Chang Eon Park, Seung-Dae Choi, YoungJae Jo, Min-Chul Kim, Min Ji Kim, Gi Ung Kang, Hyung Woo Jo, Hyunju Yun, Bora Lee, Clara Yongjoo Park, Eun Soo Kim, Sang-Jun Kim, Jae-Ho Shin

## Abstract

The gut microbiome is essential to human health. However, little is known about the influence of the environment versus host-related factors (e.g. genetic background, sex, age, and body mass) in the formation of human intestinal microflora. Here, we present evidence in support of the importance of host-related factors in the establishment and maintenance of individual gut assemblages. We collected fecal samples (n = 249) from 44 Korean naval trainees and 39 healthy people living in Korea over eight weeks and sequenced the bacterial 16S rRNA genes. The following hypotheses were tested: 1) microbiome function is linked to its diversity, community structure, and genetic host-related factors, and 2) preexisting host-related factors have a more significant effect on gut microbiome formation and composition than environmental factors. For each individual, the difference between the initial gut microbiota and that after eight weeks was negligible even though the 44 naval trainees lived in the same area and received the same diet, the same amount of exercise, and the same amount of physical stress during the study. This suggests that host-related factors, rather than environmental factors, is a key determinant of individual gut microflora. Moreover, eight weeks of physical training and experiencing the same environmental conditions resulted in an increase in the species *Bifidobacterium*, *Faecalibacterium*, and *Roseburia* in most trainees, suggesting a healthier intestinal environment.

**IMPORTANCE:** In order to understand the role of human gut microbiome, it is important to know how individual’s gut microbiota are formed. In this study, we tested the host-related factors versus environmental factors to affect gut microbiome and found that the former have a more association. However, we also found that the controlled environment give an effect on the gut microflora as well. This study provides preliminary evidence that differences in the formation and diversity of gut microbiota within a population could be determined by host-related factors rather than environmental factors.

## Introduction

The human body harbors complex and diverse microbial communities (about 10–100 trillion microbes), including bacteria, archaea, eukaryotes, and viruses. These communities may be described as the “normal flora” in healthy individuals (1–3). These organisms perform a variety of metabolic functions. For example, organisms residing in human and animal colons, with an estimated density of 10^11^–10^12^ cells/g (4–6), have been implicated in niche-specific metabolic, defense, and reproductive functions (1). They also influence their hosts during homeostasis and disease development through existing immunological factors, coexisting interacting networks of other microorganisms, and environmental influences (7).

The human gut microbiota is a reservoir of microorganisms and is crucial to health (3) through its involvement in metabolic interactions (e.g., food decomposition and nutrient intake) (8, 9), drug metabolism (10, 11), energy production and storage (12), and protection against pathogens (13). The gut microbiome provides signals that influence the development of the host immune system and stimulate the maturation of immune cells (14, 15). However, the gut microbiota is not only associated with human well-being but also with human disease conditions, including metabolic diseases, growth disorders, mental illness (such as autism), and obesity (16–20). Consequently, gut microbes affect human physiology directly and indirectly (13). Moreover, abrupt changes to the delicate balance of the microbial assemblage can result in unexpected consequences.

Despite the importance of the gut microbiome to human well-being, very little is known about the interactions between the host and the intestinal microorganisms (21). Previous studies of host interactions with intestinal microbes found that gut microorganisms play a fundamental role in nutrient metabolism by converting dietary components that escape digestion and polymers excreted by the host into easily accessible nutrients and other compounds, such as vitamins (22). Moreover, gut microbiota regulate host immune system interactions. For example, Bacteroides fragilis induces a specific IgA response that is dependent on commensal colonization factors regulation of surface capsular polysaccharides and enhances epithelial adherence (23). Considering that the in vivo production of IgA is necessary for single-strain stability, mucosal colonization, and epithelial aggregation, this bacterial influence on the host immune system is vital. Adaptive immunity has evolved to rely on an intimate association with members of the gut microbiome (23).

Due to host-bacterial mutualism, humans have not needed to evolve some metabolic pathways, such as the ability to harvest otherwise inaccessible nutrients (24). Therefore, investigating factors that influence the diversity and composition of the human gut microbiota increases our understanding of how these communities are structured, and to predict their response to environmental change. Microbial establishment in the human intestine begins at birth. Subsequently, the intestinal microflora continues to develop through successive microbial communities, until the microbial climax community colonizes the intestine. Moreover, due to co-evolutionary interactions, microbes undergo further functional modifications (24). However, the initial gut microbiome of an infant at birth is similar in composition to the mother’s (25). In addition to environmental and genetic factors (26–29), other factors, including lifestyle, diet, stress, and probiotics (30–32) have also been implicated in controlling the establishment of gut microbiota. However, it is still poorly understood whether environmental or host-related factors play a dominant role in shaping the human gut microbial community.

Most gut microbiome studies to date have focused on Westerners (mostly Europeans and North Americans); therefore, there is a gap in our knowledge about the gut microbiota of people from other parts of the world (33). In this study, we used next-generation sequencing of bacterial 16S rRNA genes to analyze the intestinal bacterial community in South Korean naval trainees during a time when the trainees were experiencing relatively similar environmental conditions.

Our first objective was to determine whether host-related factors are linked to the community structure and diversity of the intestinal microbiota of naval trainees. Given the broad range of interactions that are commonly associated with gut microbiota, we aimed to identify key functional processes. Glendinning and Free suggested that these functions could range from providing additional metabolic functions to modulating the immune system (34). Our second objective was to determine whether preexisting host-related factors have a more significant effect on the formation and composition of the intestinal microbial community than environmental factors. Turpin et al. suggested that almost one-third of fecal bacterial taxa in the intestine are heritable (35). Furthermore, ecological concepts like dispersal, species diversity and distribution, community assembly and contamination, and genetic makeup may influence the formation, development, and maintenance of the gut microbiota. Through this objective, we sought to investigate which set factors, host-related or environmental, exerts a more significant influence on the composition of the gut microbiota.

## Results

The aim of our study was to determine whether host-related background or environmental factors play a more significant role in the formation and diversity of the intestinal microbial community. Therefore, we first identified the composition of the intestinal microbial communities of the OCS trainees, who had different host-related and environmental backgrounds, on the day they were admitted (T = week 0). After living together for eight weeks at the naval OCS and being exposed to the same environmental conditions (e.g., same diet, same physical exercise, and same sleeping regimes), their intestinal microbial communities were reanalyzed at T = 4 weeks and T = 8 weeks. As a control sample, intestinal microflora samples from healthy Koreans with different host-related and environmental backgrounds were also analyzed. Theoretically, if the host-related background is more significant in determining the intestinal microbiota, the initial composition and diversity of microbial communities of naval trainees would be maintained, with negligible differences between samples collected at the beginning and end of the experimental period, and samples would remain equally distant from each other (Figure 1a). However, if environmental factors are more significant in shaping the intestinal microbiota, then the composition of microbial communities would shift, the bacterial compositions between samples would become similar, and PCoA plots would show clustering of samples (Figure 1b).

**Figure 1.**
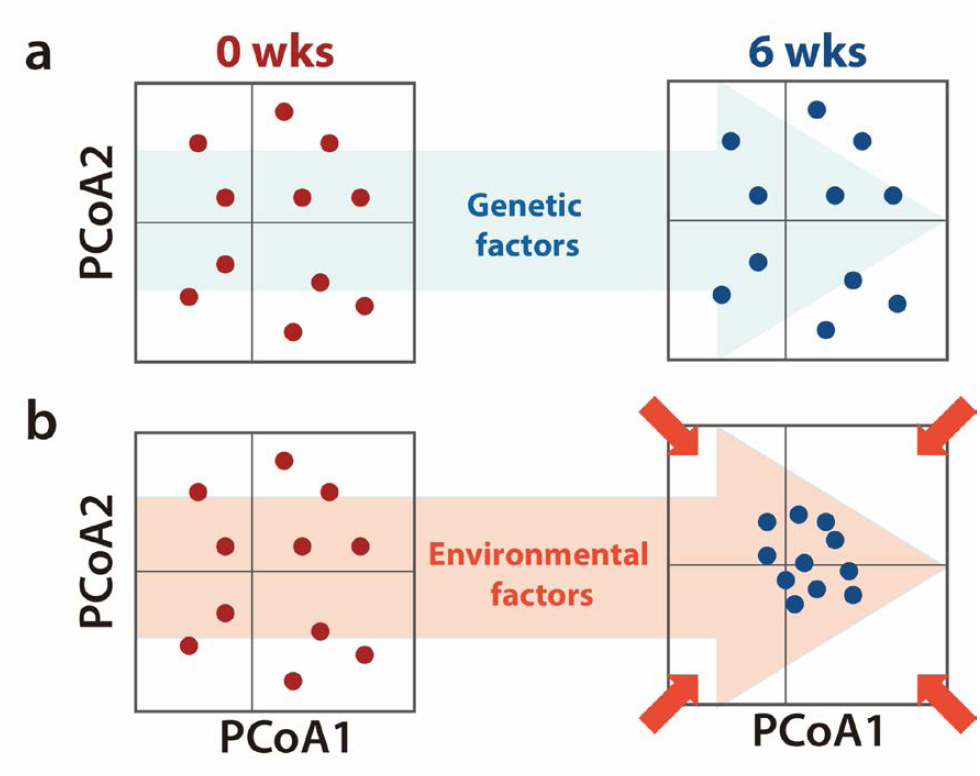
A schematic representation of how genetic and environmental factors affect the composition and structure of the host intestinal microbial community differently. (a) If genetic factors have a more significant effect, then the intestinal microbial community in each individual is maintained, and samples remain distant from each other. (b) If environmental factors have a more significant effect, then the intestinal microbial community in each individual changes, and samples will cluster together.

From 132 fecal samples collected from the 44 naval trainees (OCS), we obtained 2218 OTUs at 97% similarity from 1.49 million reads. The average number of reads per sample was 38 443 (ranging from 27 652 to 48 155). Similarly, 117 stool samples were collected from 39 healthy Korean adults as the control group. We obtained 1927 OTUs at 97% similarity from 3.19 million reads. The average number of reads per sample was 27 102 (ranging from 7331 to 45 842).

### Composition and diversity of the gut microbial community at week zero

Based on the demographic questionnaire and the FFQ, we compared characteristics of the control and experimental groups at the beginning of the experiment. We found no significant differences among most of the individuals except for age, regular exercise, recent weekly exercise time, and hours of sleep (Table 1). We then performed PCoA ordination based on weighted UniFrac distances to determine the beta diversity of the intestinal microbial communities of the experimental and control groups. The PCoA plot shows that both the experimental and control groups had distinct intestinal microbial communities, and all samples remained distant from each other (Figure 2a and 2b). The adonis function was used to determine whether host-related factors, such as sex, age, weight, height and BMI, and environmental factors, including exercise, sleep and daily nutrient intake, have a major effect on the composition of the intestinal microbial community. Adonis results show that the majority of environmental factors studied did not significantly affect the gut microbial community. Sex (P = 0.006, R2 = 0.131), height (P = 0.031, R2 = 0.085), weight (P = 0.001, R2 = 0.174), BMI (P = 0.001, R2 = 0.079), and number of bowel movements per week (P = 0.005, R2 = 0.142) show significant association with the intestinal microbiome in the control group. In the experimental group, sex (P = 0.029, R2 = 0.064), height (P = 0.008, R2 = 0.081), and weight (P = 0.007, R2 = 0.089) show significant association with the intestinal microbiome (Table 2). Alpha diversity was also calculated before admission to the navy center and was significantly higher in the experimental group than in the control group (Figure 2c–d). The Shannon index (P = 0.010, Mann Whitney test), which is the most commonly used alpha diversity index, shows an average diversity of 5.38 (3.91–7.14) for the control group, and 5.81 (3.88–6.84) for the experimental group. Phylogenetic diversity (P = 0.016, Mann Whitney test), which measures the phylogenetic relationship between OTUs, has a median value of 27.08 (15.35–43.83) for the control group, and 29.43 (10.10–37.94) for the experimental group.

**Figure 2.**
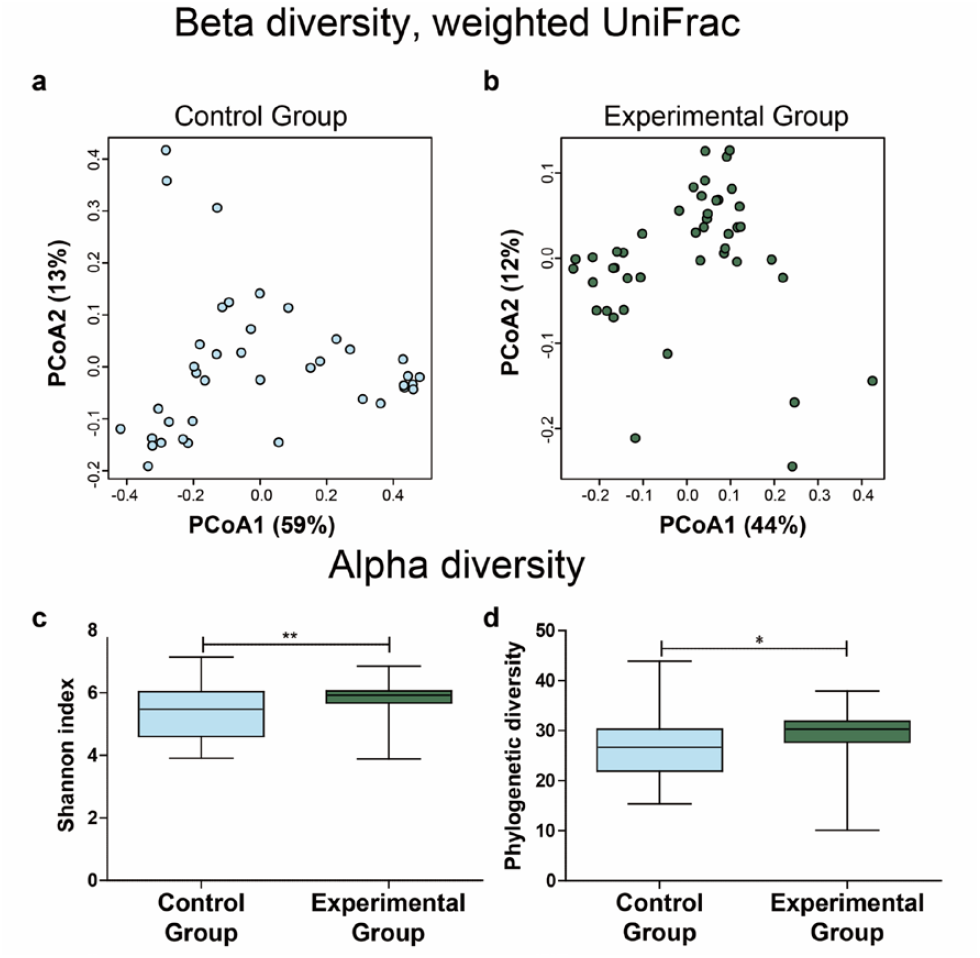
Individuals have distinctive intestinal microbial communities. PCoA plots of weighted UniFrac distances in (a) control and (b) experimental groups at T = 0 weeks. The first and second principal components (PCo1 and PCo2) are plotted. The percentage of variance in the dataset explained by each axis is reported. Variation of bacterial alpha diversity indices in intestinal microbiota between control and experimental groups: (c) Shannon index and (d) phylogenetic diversity. Data are shown as mean ± SEM; *p < 0.05, **p < 0.01 by the Mann Whitney U test.

**Table 1.** Baseline characteristics of participants in the study. Data are shown as mean ± SEM; p-values are obtained by unpaired t-test (Welch’s correction) or Mann-Whitney U test (continuous variables) or chi-square test (proportions).

| Characteristics | Control Group<br>(n = 39) | Experimental Group<br>(n = 44) | p-value |
| --- | --- | --- | --- |
| **Host-related factors** |  |  |  |
| **Sex, % (n)** |  |  | 0.056<sup>c</sup> |
| Male | 58.97 (23) | 77.27 (34) |  |
| Female | 41.02 (16) | 22.72 (10) |  |
| **Age, (years)** | 29.92 ± 1.39 | 24.25 ± 0.30 | <0.001<sup>b</sup> |
| **Height, (cm)** | 169.54 ± 1.33 | 171.00 ± 1.08 | 0.400<sup>a</sup> |
| **Weight, (kg)** | 66.21 ± 2.08 | 71.30 ± 1.71 | 0.063<sup>a</sup> |
| **Body mass index (BMI)** |  |  | 0.679<sup>c</sup> |
| <18.50 | 5.13 (2) | 2.27 (1) |  |
| 18.50-24.99 | 69.23 (27) | 65.90 (29) |  |
| 25.00-29.99 | 25.64 (10) | 31.82 (14) |  |
| >30.00 | 0.00 (0) | 0.00 (0) |  |
| **Environmental factors** |  |  |  |
| **Do you Smoke?, % (n)** |  |  | 0.924<sup>c</sup> |
| Yes | 28.20 (11) | 27.27 (12) |  |
| No | 71.79 (28) | 72.73 (32) |  |
| **Do you exercise regularly?, % (n)** |  |  | 0.019<sup>c</sup> |
| Yes | 33.33 (13) | 59.09 (26) |  |
| No | 66.66 (26) | 40.91 (18) |  |
| **If you do regular exercise, what kind of exercise do you do?, % (n)** |  |  | 0.904<sup>c</sup> |
| Weight training | 15.38 (2) | 26.92 (7) |  |
| Cardio | 15.38 (2) | 7.69 (2) |  |
| Both | 69.23 (9) | 65.38 (17) |  |
| **How much time did you exercise during the past week?, (min)** | 96.03 ± 27.00 | 208.10 ± 37.34 | 0.008<sup>b</sup> |
| **How many hours a day do you sleep?, (hr)** | 6.38 ± 0.21 | 7.69 ± 0.19 | <0.001<sup>b</sup> |
| **How is your sleep quality?, % (n)** |  |  | 0.212<sup>c</sup> |
| Very Bad | 7.69 (3) | 0.00 (0) |  |
| Bad | 10.26 (4) | 6.82 (3) |  |
| Normal | 30.77 (12) | 50.00 (22) |  |
| Good | 41.03 (16) | 31.82 (14) |  |
| Very Good | 10.26 (4) | 9.09 (4) |  |
| No Answer | 0.00 (0) | 2.27 (1) |  |
| **How many people do you live with?, (n)** | 2.69 ± 0.24 | 3.25 ± 0.21 | 0.083<sup>a</sup> |
| **Do you have pets?, % (n)** |  |  | 0.222<sup>c</sup> |
| Yes | 12.82 (5) | 22.72 (10) |  |
| No | 87.17 (34) | 75.00 (33) |  |
| No Answer | 0.00 (0) | 2.27 (1) |  |
| **How many times a week do you have a bowel movement?, (n)** | 5.49 ± 0.45 | 6.18 ± 0.24 | 0.052<sup>b</sup> |
| **How do you feel after a bowel movement?, % (n)** |  |  | 0.880<sup>c</sup> |
| Bad | 5.13 (2) | 2.27 (1) |  |
| Normal | 35.90 (14) | 40.91 (18) |  |
| Good | 48.72 (19) | 45.46 (20) |  |
| Very Good | 10.26 (4) | 11.36 (5) |  |
| No Answer | 0.00 (0) | 0.00 (0) |  |
| **Daily nutrient intake<sup>†</sup>** |  |  |  |
| Energy (kcal) | 3377.48 ± 182.33 | 3924.18 ± 254.79 | 0.089<sup>a</sup> |
| Carbohydrate (g) | 519.24 ± 29.66 | 584.84 ± 46.31 | 0.241<sup>a</sup> |
| Protein (g) | 117.02 ± 8.40 | 138.24 ± 9.821 | 0.108<sup>a</sup> |
| Fat (g) | 79.89 ± 6.48 | 95.52 ± 6.92 | 0.106<sup>a</sup> |
| Cholesterol (g) | 473.82 ± 41.44 | 603.84 ± 68.07 | 0.112<sup>a</sup> |
| Fiber (mg) | 28.11 ± 2.04 | 28.77 ± 2.09 | 0.822<sup>a</sup> |

<sup>†</sup> Daily nutrient intake were adjusted subjects number; control group (n = 28), experimental group (n = 21); FFQ non-responders were excluded, <sup>a</sup> p-values were obtained by the unpaired t-test (Welch's correction), <sup>b</sup> p-values were obtained by the Mann-Whitney U test, <sup>c</sup> p-values were obtained by the chi-square test

**Table 2.**
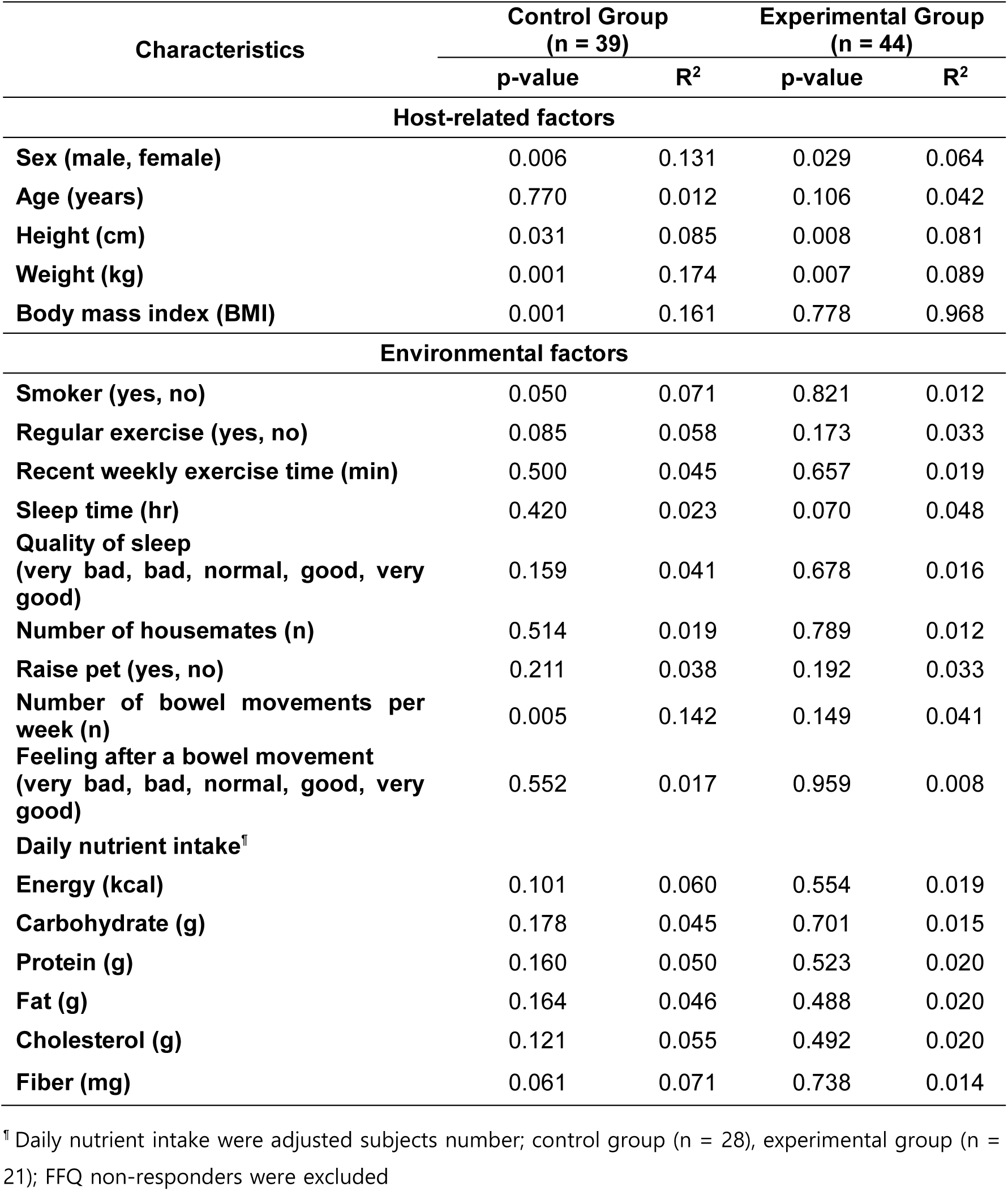
Adonis function results for host-related and environmental factors (T = 0 weeks) based on weighted UniFrac distance. Weighted UniFrac beta diversity analysis was performed using the adonis function to identify significant differences in microbiota composition.

| Characteristics | Control Group<br>(n = 39) |  | Experimental Group<br>(n = 44) |  |
| --- | --- | --- | --- | --- |
|  | p-value | R<sup>2</sup> | p-value | R<sup>2</sup> |
| **Host-related factors** |  |  |  |  |
| **Sex (male, female)** | 0.006 | 0.131 | 0.029 | 0.064 |
| **Age (years)** | 0.770 | 0.012 | 0.106 | 0.042 |
| **Height (cm)** | 0.031 | 0.085 | 0.008 | 0.081 |
| **Weight (kg)** | 0.001 | 0.174 | 0.007 | 0.089 |
| **Body mass index (BMI)** | 0.001 | 0.161 | 0.778 | 0.968 |
| **Environmental factors** |  |  |  |  |
| **Smoker (yes, no)** | 0.050 | 0.071 | 0.821 | 0.012 |
| **Regular exercise (yes, no)** | 0.085 | 0.058 | 0.173 | 0.033 |
| **Recent weekly exercise time (min)** | 0.500 | 0.045 | 0.657 | 0.019 |
| **Sleep time (hr)** | 0.420 | 0.023 | 0.070 | 0.048 |
| **Quality of sleep**<br>**(very bad, bad, normal, good, very good)** | 0.159 | 0.041 | 0.678 | 0.016 |
| **Number of housemates (n)** | 0.514 | 0.019 | 0.789 | 0.012 |
| **Raise pet (yes, no)** | 0.211 | 0.038 | 0.192 | 0.033 |
| **Number of bowel movements per week (n)** | 0.005 | 0.142 | 0.149 | 0.041 |
| **Feeling after a bowel movement**<br>**(very bad, bad, normal, good, very good)** | 0.552 | 0.017 | 0.959 | 0.008 |
| **Daily nutrient intake<sup>†</sup>** |  |  |  |  |
| **Energy (kcal)** | 0.101 | 0.060 | 0.554 | 0.019 |
| **Carbohydrate (g)** | 0.178 | 0.045 | 0.701 | 0.015 |
| **Protein (g)** | 0.160 | 0.050 | 0.523 | 0.020 |
| **Fat (g)** | 0.164 | 0.046 | 0.488 | 0.020 |
| **Cholesterol (g)** | 0.121 | 0.055 | 0.492 | 0.020 |
| **Fiber (mg)** | 0.061 | 0.071 | 0.738 | 0.014 |

<sup>†</sup> Daily nutrient intake were adjusted subjects number; control group (n = 28), experimental group (n = 21); FFQ non-responders were excluded

### Composition and diversity of gut microbial community after eight weeks of training

In the experimental group, one host-related factor and majority of environmental factors show significant differences between the first day of admission to the OCS (T = week 0), four weeks (T = 4 weeks), and eight weeks later (T = 8 weeks). The factors that differed were BMI, exercise time, the number of housemates, and daily nutrient intake (carbohydrates, cholesterol, and fiber). However, a host-related factor (BMI) and environmental factors did not statistically differ in the control group (Table 3). The intestinal bacterial communities of the experimental group did not differ among the first day of admission to the OCS (T = week 0), four weeks (T = 4 weeks), and eight weeks later (T = 8 weeks). This finding is confirmed by the PCoA plot showing dissimilarities in bacterial communities and no clustering of samples according to the measured environmental factors (Figure 3a–b). The composition of bacteria in the intestines of trainees and ordinary people remain distinct, suggesting that host-related factors have a greater effect than environmental factors in determining the composition of the intestinal microbial community. Similarly, alpha diversity indices show no significant differences at the end of the experiment in experimental and control groups (Figure 3c– f). The Shannon index shows no significant change between 0, 4, and 8 weeks (control group: F = 0.585, P = 0.559, repeated-measures one-way ANOVA; experimental group: P = 0.431, Friedman test) and neither does the phylogenetic diversity (control group: F = 0.504, P = 0.602, repeated-measures one-way ANOVA; experimental group: P = 0.084, Friedman test).

**Figure 3.**
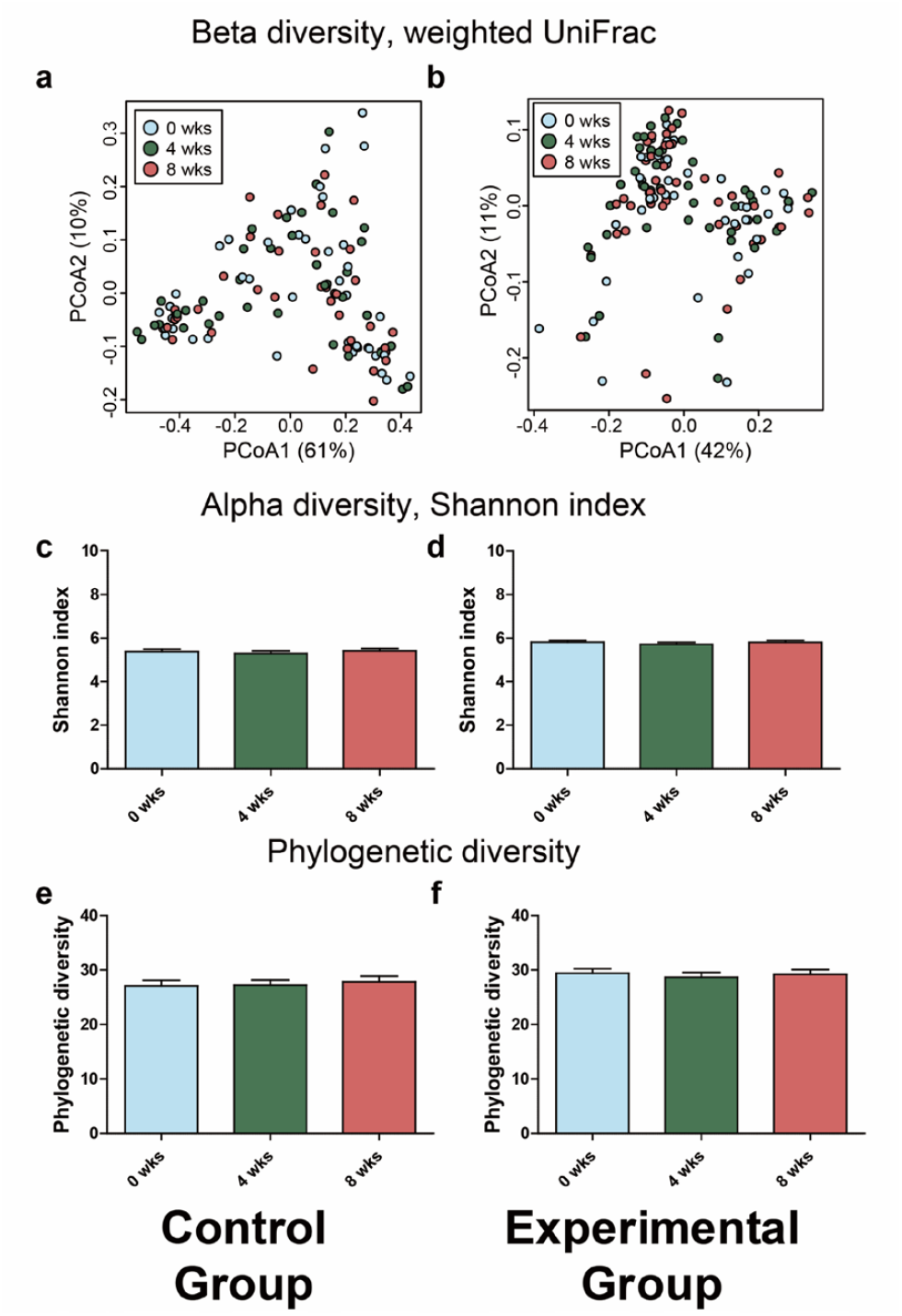
Environmental factors have negligible effects on the gut microbiome. PCoA plots of weighted UniFrac distances in (a) control and (b) experimental groups at T = 0 weeks, T = 4 weeks, and T = 8 weeks. Variation of bacterial alpha diversity indices in intestinal microbiota at T = 0 weeks, T = 4 weeks, and T = 8 weeks: Shannon index of (c) control and (d) experimental groups and phylogenetic diversity index of (e) control and (f) experimental groups. Data are shown as mean ± SEM. The alpha diversity of the control group was analyzed using repeated-measures one-way ANOVA. The alpha diversity of the experimental group was analyzed using the Friedman rank-sum test.

**Table 3.** Characteristics of participants at T = 0 weeks, T = 4 weeks, and T = 8 weeks. Data are shown as mean ± SEM; p-values are obtained by repeated-measures one-way ANOVA.

| Characteristics | Control Group (n = 39) |  |  |  | Experimental group (n = 44) |  |  |  |
| --- | --- | --- | --- | --- | --- | --- | --- | --- |
|  | 0 wks | 4 wks | 8 wks | p-value | 0 wks | 4 wks | 8 wks | p-value |
| Body mass index (BMI) | 22.83 ± 0.48 | 22.73 ± 0.46 | 22.77 ± 0.47 | 0.996 | 24.22 ± 0.41 | 23.96 ± 0.38 | 23.25 ± 0.31 | <0.001 |
| Recent weekly exercise time (min) | 88.33 ± 27.35 | 99.87 ± 25.52 | 89.23 ± 24.81 | 0.734 | 208.10 ± 37.34 | 3252.00 ± 32.35 | 1311.00 ± 24.43 | <0.001 |
| Sleep time (hr) | 6.38 ± 0.21 | 6.51 ± 0.19 | 6.44 ± 0.19 | 0.835 | 7.69 ± 0.19 | 7.51 ± 0.10 | 7.63 ± 0.09 | 0.552 |
| Number of housemates (n) | 2.69 ± 0.24 | 2.39 ± 0.23 | 2.38 ± 0.22 | 0.543 | 3.25 ± 0.21 | 4.00 ± 0.00 | 4.00 ± 0.00 | <0.001 |
| Number of bowel movements per week (n) | 25.44 ± 0.46 | 6.08 ± 0.56 | 6.09 ± 0.61 | 0.865 | 6.17 ± 0.24 | 5.55 ± 0.44 | 5.36 ± 0.28 | 0.083 |
| Daily nutrient intake<sup>1</sup> |  |  |  |  |  |  |  |  |
| Energy (kcal) | 3377.48 ± 182.33 | 3330.89 ± 182.47 | 3284.30 ± 189.63 | 0.209 | 3924.18 ± 254.79 | 3474.01 ± 11.03 | 3369.69 ± 24.78 | 0.055 |
| Carbohydrate (g) | 519.24 ± 29.66 | 512.84 ± 29.07 | 506.43 ± 30.12 | 0.367 | 584.84 ± 46.31 | 468.66 ± 1.08 | 464.84 ± 3.92 | 0.017 |
| Protein (g) | 117.02 ± 8.40 | 116.24 ± 8.43 | 115.45 ± 8.54 | 0.332 | 138.24 ± 9.82 | 139.70 ± 0.88 | 135.12 ± 1.36 | 0.694 |
| Fat (g) | 79.89 ± 6.48 | 78.08 ± 6.42 | 76.28 ± 6.53 | 0.091 | 95.52 ± 6.92 | 112.22 ± 0.54 | 104.29 ± 0.62 | 0.051 |
| Cholesterol (g) | 473.82 ± 41.44 | 466.13 ± 41.17 | 458.44 ± 41.25 | 0.052 | 603.84 ± 68.07 | 836.35 ± 11.31 | 733.68 ± 21.46 | 0.006 |
| Fiber (mg) | 28.11 ± 2.04 | 27.69 ± 2.02 | 27.26 ± 2.03 | 0.095 | 28.77 ± 2.09 | 37.99 ± 0.57 | 39.81 ± 0.62 | <0.001 |

<sup>1</sup> Daily nutrient intake were adjusted subjects number; control group (n = 28), experimental group (n = 21); FFQ non-responders were excluded

Changes in the intestinal microbiota for each experimental period were measured using the distance matrix (Figure 4).

**Figure 4.**
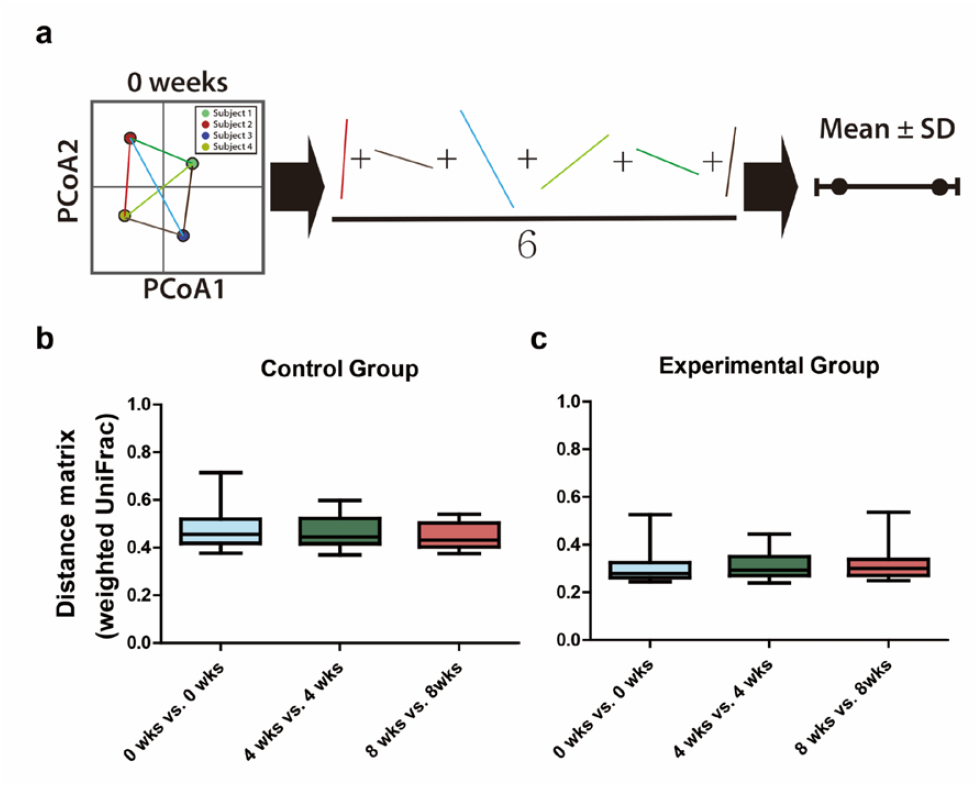
Even when environmental factors are controlled, individual intestinal microbial communities are maintained. (a) A schematic representation of the average weighted UniFrac distance on the composition of the intestinal microbial community. The average distance between all samples was used to determine how differentiated the intestinal microflora of subjects were at each time point. The average weighted UniFrac distances among (b) control and (c) experimental groups within each time point are shown in the box plot. Data were analyzed using the Kruskal-Wallis test.

This study was designed to determine the distribution of the intestinal microbial community for each subject at each sampling time. The average distance between all samples was used to determine the degree to which the intestinal microflora of subjects differed at each time point (Figure 4a).

For the control group (Figure 4b), there is no significant change (P = 0.204, Kruskal-Wallis test) between T = 0 weeks (mean = 0.47, 0.38–0.71), T = 4 weeks (mean = 0.47, 0.37–0.60), and T = 8 weeks (mean = 0.45, 0.38–0.54).

The experimental group (Figure 4c) also shows no significant change (P = 0.238, Kruskal-Wallis test) between T = 0 weeks (mean = 0.30, 0.24–0.53), T = 4 weeks (mean = 0.31, 0.24–0.44), and T = 8 weeks (mean = 0.32, 0.25–0.53). Despite highly controlled environmental factors, the composition of the intestinal microbial community does not seem to be affected.

The distance matrix was again used to illustrate changes in the intestinal microbial community (Figure 5).

**Figure 5.**
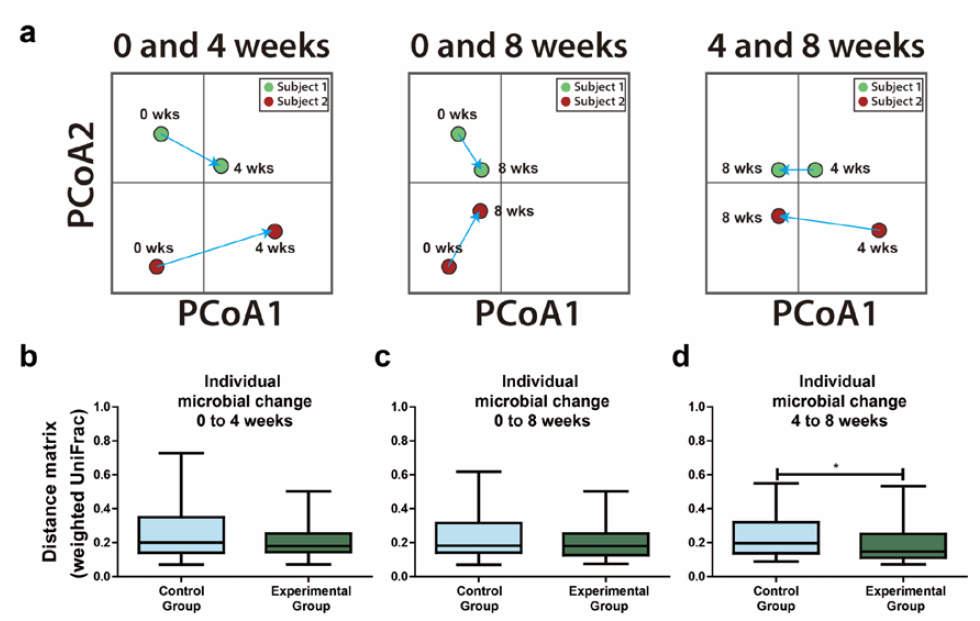
Regular living stabilizes the gut microbiome in a healthy direction. (a) A schematic representation of the variation in individual weighted UniFrac distances according to sampling time, explaining how individual intestinal microflora changes over time. Changes in gut microbiome between (b) T = 0 weeks and T = 4 weeks, (c) T = 0 weeks and T = 8 weeks, and (d) T = 4 weeks and T = 8 weeks of the control and experimental groups. *p < 0.05 by the Mann Whitney U test.

Changes in intestinal microflora over time were tracked for each subject (Figure 5a). The results showed that there is no significant difference (P = 0.154) in the individual distance matrix between T = 0 weeks and T = 4 weeks. The mean distance in the control group is 0.26 (ranging between 0.07 and 0.73), and in the experimental group, the mean distance measures 0.20 (ranging between 0.07 and 0.50), as shown in Figure 5b. Similarly, the individual distance matrices do not differ between the control group (mean = 0.24; 0.07–0.31) and the experimental group (mean = 0.20; 0.08–0.50) between T = 0 weeks and T = 8 weeks (P = 0.516), as shown in Figure 5c. However, between T = 4 weeks and T = 8 weeks, the individual distance matrices are significantly different (P = 0.033) between the control group (mean = 0.23; 0.09–0.55) and the experimental group (mean = 0.19; 0.07–0.53), as shown in Figure 5d. This suggests that environmental factors do not exert major effects on the composition of the intestinal microbial community but they do have some influence.

### Shifts in intestinal microbial communities after eight weeks

The majority of bacterial sequences recovered in our study belonged to the following phyla: Firmicutes (mean = 5413.70, 3104–6925) and Bacteroidetes (mean = 2319.19, 219–3343), which do not differ significantly between T = 0 weeks and T = 8 weeks.

The relative abundance of the top five groups (at the phyla, class, order, and family levels) was compared between T = 0 weeks and T = 8 weeks in the experimental group (Figure S1). The class Actinobacteria (P = 0.022), the order Bifidobacteriales (P = 0.026), and the families Bifidobacteriaceae (P = 0.026) and Ruminococcaceae (P = 0.019) increase significantly at T = 8 weeks. However, the phylum Proteobacteria (P = 0.003), the class Coriobacteriia (P = 0.0001), the order Coriobacteriales (P = 0.0001), and the family Lachnospiraceae (P = 0.042) decrease significantly at T = 8 weeks. At the genus level, *Bifidobacterium* sp. (P = 0.026), *Faecalibacterium* sp. (P = 0.045), and *Roseburia* sp. (P = 0.020) significantly increase at T = 8 weeks. However, *Blautia* sp. (P = 0.015), *Collinsella* sp. (P = 0.0001), and *Ruminococcus* sp. (P = 0.022) decrease significantly at T = 8 weeks (Figure 6). In the control group, intestinal microorganisms do not change significantly, unlike the experimental group (Figure S2).

**Figure 6.**
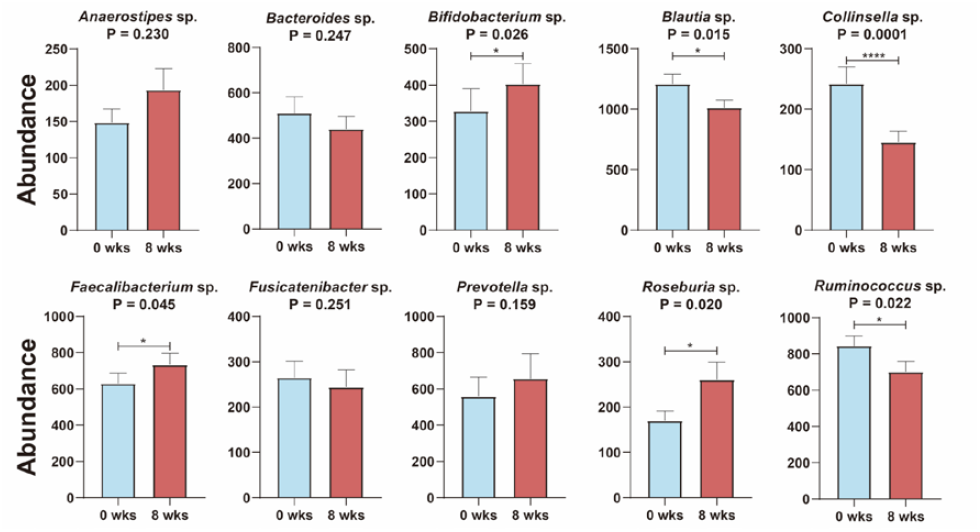
Controlled living conditions of the OCS training cause beneficial changes to the gut microbial community. Abundance of intestinal bacteria by family. Data are shown as mean ± SEM. *p < 0.05, **p < 0.01, ***p < 0.001, ****p < 0.0001. *Blautia*, *Fusicatenibacter*, and *Ruminococcus* were analyzed using a paired t-test. *Anaerostipes*, *Bacteroides*, *Bifidobacterium*, *Collinsella*, *Faecalibacterium*, *Prevotella*, and *Roseburia* were using the Wilcoxon matched-pairs signed-rank test.

## Discussion

The primary goal of microbiome research is to elucidate the factors that determine microbial structure and composition within and upon a host (36). To this end, the current study demonstrated that variations in the formation and composition of the human intestinal microbiota are primarily determined by host-related factors rather than environmental factors. Previous studies (37, 38) support our findings. Moreover, the results of Kolde et al. suggest that host genetic factors are vital in maintaining gut microbiome interactions (39). Furthermore, it has been proposed that the genetic background of the host may influence the composition and formation of the intestinal microbial community.

One host-related factor (age) and some environmental factors, including regular exercise and exercise time, and sleep time, showed significant differences between the experimental and control groups (Table 1). In the case of the experimental group, we propose that these differences are because the subjects were enlisted as military officer candidates. Alpha diversity was significantly higher in the experimental group than in the control group (Figure 2c–d). This may be due to differences in specific environmental factors in the experimental group. According to Clarke et al. (40), athletes frequently have high intestinal microbial diversity. In addition, we found that similarity in intestinal microbial communities occurs more frequently when specific host-related factors (sex, height, weight, BMI) and an environmental factor (the number of bowel movements per week) are similar (Table 2). Our results mirror the findings of Oki et al. (41), and Haro et al. (42), who report that host-related and environmental factors have a notable effect on the formation of the intestinal microbiome in various cohorts. Compared with the control group, the experimental group was not significantly different in BMI and the number of bowel movements per week. This is probably because the majority of individuals in both groups had similar values for these factors.

### Hypothesis 1: The structure and diversity of the gut microbial community are linked to genetic host-related factors

The results reported in this study (Figure 3) support our hypothesis that host-related factors control the host gut microflora. Our results also showed that host-related factors had a considerable effect on the composition of the intestinal microbial community. This is shown in the PCoA plot, which clusters intestinal bacterial communities by individual subjects under a controlled environment. The results showed that samples remained distinct from each other after trainees shared identical environmental conditions for eight weeks (Figure 3b). Bacterial diversity showed the same pattern as community composition, with no significant changes at the end of the experimental period (Figure 3b). Together, these findings suggest that genetically-driven intrinsic factors exist that may organize the gut microbiota structure and participate in the formation of the microbial community. Although the cause-and-effect relationships behind these links have yet to be elucidated, host genetics does have an impact on host health (43). Although our results showed that the environmental influence on the gut microbiota in our cohort is limited, we suggest that greater interaction between genetic and environmental factors exists, which may contribute to the genetic influence observed in this study. Small et al. investigated the importance of genotype-environment interactions in understanding host genetic mechanisms and complex molecular relationships between hosts and their resident microbes (44). Similar studies using genetic techniques have demonstrated that the gut microbiota is influenced by both environmental and genetic factors (29, 45, 46). Genetic factors have been proposed to control the composition and diversity of gut microbiota under controlled environmental conditions. However, genetics cannot be the only factor that influences the intestinal microbial community because the intestine can only be colonized by microorganisms present in the environment (47).

Our findings are also consistent with a study by Org et al. (48). They reported that genetic background, under controlled environmental conditions, plays a considerable role in the composition and diversity of the intestinal microbial community. Host genes that are associated with the gut microbiota regulate diet-sensing, metabolism, and host immunity (49). These background interactions may, in turn, influence the nature and structure of the overall host gut microbiota. Other studies have demonstrated that the host genetic background has a significant effect on the composition of the gut microbial community. Henao-Mejia et al. (50) and Peng et al. (51) showed that mice with diabetes or mutations in inflammatory signaling genes differ in their gut microbial composition relative to wild-type mice. Functionalities associated with the genome encoded by the gut microbiome assemblage expand the host’s physiological potential by increasing digestive capabilities, priming the immune system, producing vitamins, degrading xenobiotics, and resisting colonization by pathogens (49). Governing these host gene-gut microbiota interactions are ecological and evolutionary consequences that often emerge from complex interspecies relationships (44). In a related finding, Buhnik-Rosenblau et al. (52) reported that because the gut microbiota is strongly associated with the health status of the host, its composition may be affected by environmental factors, such as diet and maternal inoculation. Moreover, the operational functions behind the host’s genetic control over gut microbiota are consistent with the broader effects of evolutionary divergence of the gut microbiota composition (29).

### Hypothesis 2: Preexisting host-related factors have a more significant effect than environmental factors on the formation and composition of the intestinal microbial community

Our hypothesis was again supported by our results because the abundance of certain types of microbial taxa was consistently and significantly associated with host-related factors. The relative abundance of the two most abundant microbial taxa (i.e., Bacteroidetes and Firmicutes) showed slight differences at the beginning and end of the experiment, with variation between study groups (Figure S1–S2). This finding is consistent with that reported by Richards et al. (53) when they sought to identify host genes responsible for microbiome regulation. Koliada et al. (54) also found that Bacteroidetes and Firmicutes are abundant in the gut microbiota of healthy obese individuals in Ukraine. However, in the present study, no enrolled subjects were obese at enrollment and did not have a history of obesity. This agrees with the findings of Clarke et al. (55) that Bacteroidetes and Firmicutes are found in both lean and obese individuals in varying proportions. However, the relative abundance of these taxa in the present study was not related to lean or obese status. Correlations between the relative abundance of gut microbes and genetic loci have been found in mice following the same diet (29, 45) and in humans with Crohn’s disease (27). It is likely that variations in the gut microbiome are governed by molecular mechanisms, such as changes in gene regulation in the host epithelial cells that directly interface with the gut microbiota (53).

### Environmental factors also affect intestinal microbial community formation

Controlling for environmental factors, such as regular sleep, a well-balanced diet, and steady exercise, the genera *Bifidobacterium*, *Faecalibacterium*, and *Roseburia* significantly increased in abundance by the end of the experimental period (Figure 6). *Bifidobacterium* is a significant component of gut microflora and plays a crucial role in human health (56). Furthermore, *Bifidobacterium* regulates intestinal homeostasis, modulates local and systemic immune responses, and protects against inflammatory and infectious diseases (57, 58). The abundance of *Bifidobacterium* was found to significantly increase in rats undergoing moderate exercise (59). *Bifidobacterium* is also associated with a healthier status in adults; Aizawa et al. (60) reported a decrease in *Bifidobacterium* in patients with severe depression relative to people without severe depression. Similarly, *Faecalibacterium*, which promotes a healthy digestive tract by producing butyrate and lowering the oxygen tension of the lumen (61), significantly increased in our experiment. These results mirror the findings of Campbell et al. (62), who reported an increase in *Faecalibacterium prausnitzi* in rats after physical exercise and concluded that *F. prausnitzi* might protect the intestine through oxygen detoxification by a flavin/thiol electron shuttle. *Roseburia* also plays a role in maintaining intestinal health and immune defense systems (e.g., regulatory T cell homeostasis) through the production of butyrate (63). The abundance of *Roseburia* decreases in many intestinal diseases and has been used as an indicator of intestinal health (64). These bacteria ferment insoluble fiber as an energy source (65). The abundance of *Roseburia* decreases as the host intake of carbohydrate decreases (66) or fat increases (67). In our results, carbohydrate intake significantly decreased after training, but the abundance of *Roseburia* showed an opposite pattern. We propose that the increased fiber intake during training produced these results. *Faecalibacterium prausnitzii* and *Roseburia* are known as key organisms in the formation of a healthy microbiota (19, 68).

Some taxa decrease significantly during training. For example, the abundance of Proteobacteria had significantly decreased by the end of the study. Bacteria belonging to the Proteobacteria phylum cause dysbiosis in the intestinal microflora (69). They are found in many patients with irritable bowel disease and are known to cause inflammation (70). Inflammation increases oxygen levels in the large intestine, which reduces the absolute anaerobic bacteria that constitute most of the intestinal microflora, resulting in dysbiosis (71). Our results indicated that, due to the controlled living conditions imposed during training, the abundance of microorganisms that produce short-chain fatty acids increased. Therefore, the internal anaerobic condition was maintained, resulting in a decrease in Proteobacteria.

The abundance of bacteria of the genera *Collinsella* and *Ruminococcus* decreased by the end of the study. *Collinsella* was increased in obese pregnant women with low fiber diet group (72). It is also reported that this genus increases as carbohydrate intake decreases (73). Diet changes during training could explain the decrease in *Collinsella*. *Ruminococcus* showed the same pattern as *Collinsella* because *Ruminococcus gnavus* and *R. torques* decreased significantly during training (Figure S3). This result suggests that the controlled living routine of the OCS training increases the abundance of bacteria that have a beneficial effect on the host and reduces the abundance of bacteria that have a harmful effect. *Ruminococcus* is an enterotype that enters the intestine (74) and has the ability to ferment complex carbohydrates, such as cellulose, pectine, and starch (73, 75), and is a producer of acetate and propionate (76, 77). The *Ruminococcus* genus is quite heterogeneous, including both beneficial and harmful species. For example, *Ruminococcus bromii* is known to exert beneficial effects on health (77), whereas other *Ruminococcus* species are proinflammatory (78, 79). Recently *Ruminococcus gnavus* and *R. torques* are reported to be associated with allergic diseases, Crohn’s disease in infants, and autism spectrum disorders (80–82). Overall, the increases in *Bifidobacterium*, *Faecalibacterium*, and *Roseburia* observed in our study indicate an improvement in the health of the intestinal environment. The effects of dietary changes on gut metabolism due to the strict naval training regime that the subjects of this study followed may have caused the proliferation of these bacterial taxa. This has previously been noted by Thomas et al. (83), who investigated the effects of inter-individual microbiota differences, focusing on the presence or absence of keystone species involved in butyrate metabolism.

Further studies in large population-based cohorts are necessary to gain a greater understanding of the relationship between the host genetic profile, gut microbiome composition, and host health (39, 43). Moreover, it will be important to map the loci that control microbiota composition and prioritize the investigation of candidate genes to improve our understanding of host-microbiota interactions (48).

The present work is a preliminary study showing that differences in the formation and diversity of intestinal microbial communities within a population are primarily determined by host-related factors rather than environmental factors. Long-term experiments, more controlled environmental conditions, and more detailed metadata are required to elucidate the factors that control the function of the gut microbiota. However, it is challenging to standardize human environmental conditions and to ignore the influence of unique lifestyle factors on a community. The results reported here support findings from recent studies (29, 36, 44, 48) suggesting that the microbiome depends more on host-related factors and less on related environmental factors for its variation, composition, and control. Finally, according to Kolde et al. (39) and Richards et al. (53), manipulating the microbiome alter the expression of the host genes. Therefore, knowledge of healthy microbiota for each genetic types suggests future therapeutic routes for human wellness.

## Materials and Methods

### Sampling

The present study was approved by the Institutional Review Board of Kyungpook National University (KNU 2017-84), and Armed Forces Medical Research Ethics Review Committee (AFMC-17-IRB-092), Republic of Korea. All subjects gave written informed consent in accordance with the Declaration of Helsinki.

Fecal samples were collected from 44 trainees of the naval officer candidate school (OCS) on the first day of enlistment (week 0) and at four and eight weeks after their admission to the naval center. The trainees lived in the same environment for eight weeks, ate the same food at regular intervals, and participated in similar training and sleeping regimes. Fecal samples were also collected from 39 healthy people living in Korea at the same sampling points as the OCS as controls. All samples were collected by participants using Transwab tubes (Sigma, Dorset, UK) and sent to the laboratory, where they were stored at −80 °C until DNA extraction.

### Data Collection

Participants were asked to complete a self-administered questionnaire to collect demographic, lifestyle, and physical activity data at weeks 0, 4, and 8. Dietary consumption was assessed by a food frequency questionnaire (FFQ) used in the 2017 Korea National Health and Nutrition Examination Survey conducted by the Korea Centers for Disease Control and Prevention. The FFQ was completed by the control and the experimental groups at week 0 and in the experimental group at weeks 4 and 8. Reported intakes below 500 kcal/d or >5,000 kcal/d for controls and >6,000 kcal/d for trainees were determined inaccurate and excluded from further analyses. The OCS provided naval trainees’ menus which were analyzed for nutrient content using the computer-aided nutritional analysis program (CAN Pro 5.0, Korea Nutrition Society). The daily nutrient intake for foods consumed by trainees that were not available in CAN Pro 5.0 were calculated by referencing the Korean Food Composition Database, Version (2019, Rural Development Administration). Meals were served buffet style, thus the analyzed nutrients were based on the ideal diet intake for trainees. Five day menus immediately prior to each naval trainees’ fecal collection date were used to estimate mean daily nutrient intake for weeks 4 and 8 of the naval trainees.

### DNA extraction, PCR amplification, and sequencing

Genomic DNA was extracted from approximately 500 μL (wet weight) of each sample using QIAamp PowerFecal DNA Isolation kits (Qiagen, Hilden, Germany), following the manufacturer’s instructions. Extracted DNA was assessed for quality by electrophoresis and was quantified using a Qubit 2.0 Fluorometer (Life Technologies, Carlsbad, CA, USA). DNA isolated from each sample was amplified using the universal primers, 515 F (5′-barcode-GTGCCAGCMGCCGCGGTAA-3′) and 907 R (5′-barcode-CCGYCAATTCMTTTRAGTTT-3′), targeting the V4-V5 regions of prokaryotic 16S rRNA genes. The barcode is an eight-base sequence unique to each sample. PCR experiments were performed under the following conditions: 95 °C for 5 min, 5 cycles of 57 °C for 30 s and 72 °C for 30 s, and 25 cycles of 95 °C for 30 s and 72 °C for 30 s. PCR was performed in duplicate in 24 μL reaction volumes, consisting of 20 μL Emerald AMP GT PCR 1X Master Mix (Takara Bio, Shiga, Japan), 0.5 μL (10 μM) of each barcoded PCR primer pair, and 3 μL of DNA template (10–50 ng DNA). PCR products were purified using an AMPure XP bead purification kit (Beckman Coulter, Brea, CA, USA) and pooled in equal concentrations. An Agilent 2100 Bioanalyzer (Agilent Technologies, Santa Clara, CA, USA) was used to confirm the correct concentration needed for sequencing. Each amplified region was sequenced on an Illumina MiSeq sequencing platform (Illumina, San Diego, CA, USA) using a MiSeq Reagent Kit v3 (Illumina, San Diego, CA, USA), according to the manufacturer’s protocols.

### Sequence processing

Raw FASTQ files were processed using QIIME version 1.9.1^84^ and quality filtered with Trimmomatic (85) using the following criteria: MINLEN:250 CROP:250. After chimera removal, all sequences were clustered into operational taxonomic units (OTUs) at 97% identity and classified against the SILVA database (v132) for 16S rRNA genes using the VSEARCH pipeline. To correct for differences in the number of reads, which can bias downstream analyses estimates, all samples were rarefied at an even sequencing depth of 7,331 reads per sample.

### Statistical analysis

Alpha and beta diversities and phylogenetic diversity were calculated in QIIME. PCoA plots were generated using the weighted UniFrac distance to visualize the beta diversity. CALYPSO software was used for the interpretation and comparison of taxonomic information from 16S rDNA datasets (Davenport, 2016). The D’Agostino-Pearson Omnibus test was used to determine the distribution of data in RStudio 1.0.153 (https://www.rstudio.com/).

The Pearson chi-square test or Fisher’s exact test was used to compare categorical variables. Statistical analysis was performed by repeated-measures one-way analysis of variance (ANOVA), ordinary one-way ANOVA, Friedman, and Kruskal-Wallis tests for multiple comparisons. Paired or unpaired Student’s t-tests with Welch’s correction were used to analyze normal data, and a Wilcoxon matched-pairs signed-rank test or Mann-Whitney U-test was used to analyze non-normal data. A p-value of < 0.05 was considered significant. Analyses were performed using Prism 8 software (GraphPad Software, San Diego, CA).

Data were tested to determine whether diversity indices and relative abundances of taxonomical groups were significantly different between samples collected at different time points. Variations in the composition of microbial communities were analyzed using the adonis function (a nonparametric method that is analogous to ANOVA) with a weighted UniFrac distance and 999 permutations. All analyses were performed using the “vegan” package in RStudio 1.0.153 (https://www.rstudio.com/).

### Data Availability Statement

The datasets generated during and/or analyzed during the current study are available from the NCBI Sequence Read Archive database under accession numbers PRJNA596059 (Experimental group) and PRJNA596112 (Control group).

### Ethics Statement

The present work was approved by the Institutional Review Board of Kyungpook National University, South Korea (KNU 2017-84), and by the Korea Armed Forces Medical Research Ethics Review Committee (AFMC-17-IRB-092).

## Acknowledgments

We thank the Republic of Korea Naval Academy (ROKNA) for their cooperation.

The study was supported by the National Research Foundation of Korea (NRF-2018R1D1A1B07044147) and the Strategic Initiative for Microbiomes in Agriculture and Food funded by Ministry of Agriculture, Food and Rural Affairs (918010–4), Republic of Korea.

No potential conflicts of interest were disclosed.

---

**Figure S1.**
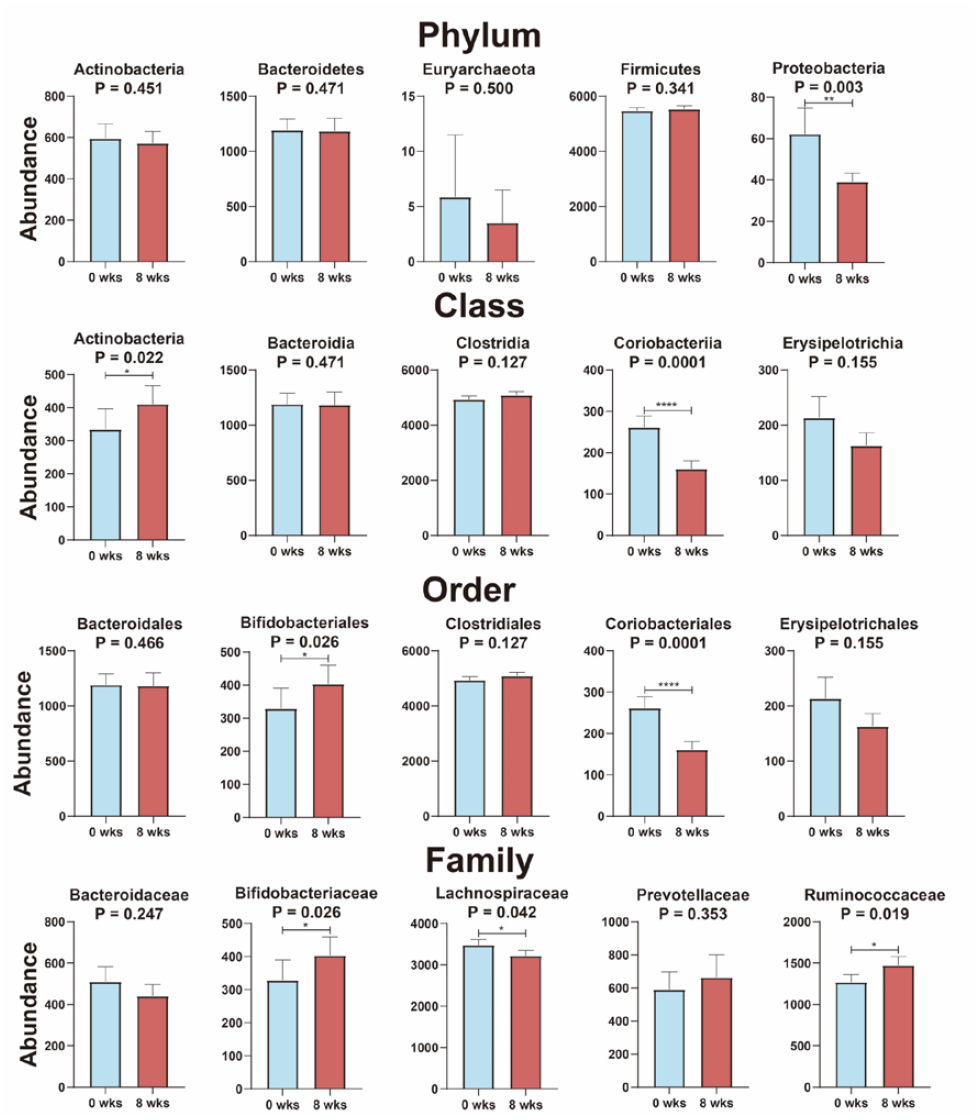
Comparison of the abundance of the detected phyla, classes, orders and families in the intestine of experimental gorup at T = 0 week and T = 8 weeks. Data are shown as mean ± SEM. *p < 0.05, **p < 0.01, ***p < 0.001, ****p < 0.0001. Bacteroidetes, Firmicutes, Bacteroidia, Clostridia, Bacteroidales, Clostridiales, Ruminococcaceae were using paired t-test. Actinobacteria (phylum), Euryarchaeota, Proteobacteri, Actinobacteria (class), Coriobacteriia, Erysipelotrichia, Bifidobacteriales, Coriobacteriales, Erysipelotrichales, Bacteroidaceae, Bifidobacteriaceae, Lachnospiraceae and Prevotellaceae were using Wilcoxon matched-pairs signed-rank test.

**Figure S2.**
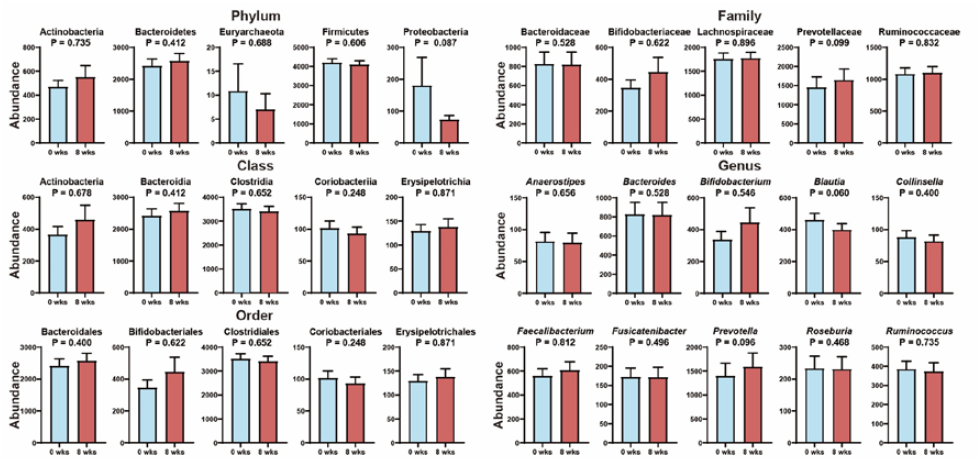
Comparison of the abundance of the detected phyla, classes, orders, families and genera in the intestine of control group at T = 0 week and T = 8 weeks. Data are shown as mean ± SEM. Bacteroidetes, Firmicutes, Bacteroidia, Clostridia, Bacteroidales, Clostridiales, Lachnospiraceae and Ruminococcaceae were using paired t-test. Actinobacteria (phylum), Euryarchaeota, Proteobacteri, Actinobacteria (class), Coriobacteriia, Erysipelotrichia, Bifidobacteriales, Coriobacteriales, Erysipelotrichales, Bacteroidaceae, Bifidobacteriaceae, Prevotellaceae, *Anaerostipes*, *Bacteroides*, *Bifidobacterium*, *Blautia, Collinsella*, *Faecalibacterium*, *Fusicatenibacter, Prevotella, Roseburia* and *Ruminococcus*.

**Figure S3.**
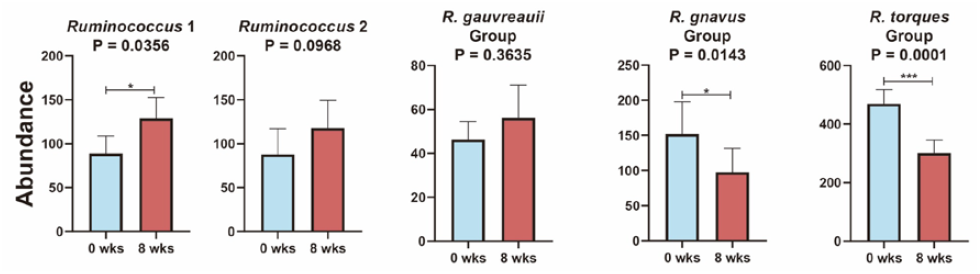
Comparison of gut microbiota within the genus *Prevotella* at T = 0 week and T = 8 weeks. *p < 0.05, **p < 0.01, ***p < 0.001 by Willcoxon matched-pairs signed-rank test.

